# Environmental luminance impairs socio-sexual recognition memory through a succinct retina to supraoptic nucleus circuit

**DOI:** 10.1101/2022.11.08.515735

**Authors:** Yu-Fan Huang, Po-Yu Liao, Jo-Hsien Yu, Shih-Kuo Chen

## Abstract

Social memory between the same gender or even different gender is a complex and heavily modulated process in the nervous system. It is important for an individual to form social memory between the opposite sex to either increase mating opportunities with multiple partners or form monogamous pair bonding. Therefore, a specific neuronal circuit to regulate social sexual memory may enhance the mating opportunity for an individual. It has been shown that both the auditory and somatosensory systems could increase the activity of oxytocin neurons in the paraventricular nucleus to regulate social behaviors. Although light exposure could influence various forms of memory, such as fear and object memory, how luminance signals modulate social recognition memory remains unclear. Here, we show that acute light exposure could impair the socio-sexual recognition memory (SSRM) in male mice. Contrary to sound and touch, light stimulation could inhibit oxytocin neurons in the SON (SON^OT^) through M1 SON-projecting ipRGCs and GABAergic neurons in the peri-SON (pSON^GABA^). Optogenetic activation of SON^OT^ neurons with channelrhodopsin is sufficient to enhance the SSRM performance in male mice, even under light conditions. Our results show that the visual system could modulate SSRM through a succinct ipRGCs-pSONGABA-SONOT neuronal circuitry. Together, we demonstrate a dedicated neuronal circuit of how luminance affects memory formation for an individual toward different sex through the oxytocin system, a powerful modulatory neurohormone in the central nervous system.

## Introduction

Oxytocin signaling is crucial for normal social/sexual behavior and recognition memory performance (Ferguson et al., 2000; Lukas et al., 2011; Nakajima et al., 2014; Oettl et al., 2016; Ross & Young, 2009). Reduction of oxytocin neurons number has been shown to be associated with autism spectrum disorder. The pair bonding and parental behaviors are also highly regulated by oxytocin and its receptor in the brain. The supraoptic nucleus (SON) and the paraventricular nucleus (PVN) in the hypothalamus (Armstrong, 2015; Liao et al., 2020) are two major regions that contain oxytocin-expressing neurons. Oxytocin neurons in those two regions could modulate neural activity in the brain through their central projections (Liao, 2021) and somatodendritic release (Ludwig, 1998). Many brain regions involved in memory formation and social interaction, including the hippocampus, central amygdala, medial amygdala, periaqueductal gray (PAG), and nucleus accumbens (NAc), are innervated by the central projection of oxytocin neurons or express oxytocin receptors (Eliava et al., 2016; Liao et al., 2020; Xiao et al., 2017). Therefore, identification of the upstream circuit which modulates oxytocin neurons is important for the understanding of social behavior regulation in the brain. Recent studies showed that signals from the sensory system could influence the activity of oxytocin neurons in the PVN. For example, physical contact such as soft touch from the somatosensory system during social interaction (Resendez et al., 2020; Tang et al., 2020; Yu et al., 2022) could activate oxytocin neurons in the PVN. Pup’s calling during parenting from the auditory system could also increase the oxytocin neuron activity in the PVN (Carcea et al., 2021). However, whether other types of sensory information such as the visual system could directly modulate the activity of oxytocin neurons in the PVN or the SON remains unclear.

In addition to image-forming functions, animals’ visual system also conveys light information to influence many physiological processes and behaviors. Recent studies suggest that light has acute effects on physiology and cognitive functions, such as sleep (Chellappa et al., 2013; Lupi et al., 2008; Zhang et al., 2021), alertness/arousal (Badia et al., 1991; Cajochen et al., 2000), anxiety (Hughes et al., 2014; Valle, 1970), and mood (LeGates et al., 2012). Furthermore, it has been shown that acute light exposure in mice could impair object and odor recognition memory (Hasan et al., 2021; Tam et al., 2016), which suggests that complex cognitive functions such as learning and memory could also be influenced by environmental luminance. However, whether there is a neuronal circuitry within visual system that could directly contribute to social/sexual recognition memory regulation remains unclear. Intrinsically photosensitive retinal ganglion cells (ipRGCs) make up most of the retinal innervation to many brain regions for non-image-forming functions including suprachiasmatic nucleus (SCN), olivary pretectal nucleus (OPN), peri-lateral habenula (pLH), intergeniculate leaflet (IGL), and supraoptic nucleus (SON) (Hattar et al., 2006). By expressing the photopigment melanopsin, these ipRGCs could directly sense light and mediate the acute effects of light on physiological functions such as arousal/anxiety (Milosavljevic et al., 2016), body temperature, sleep (Rupp et al., 2019), mood (Fernandez et al., 2018), pupillary light reflex (Chen et al., 2011), and circadian photoentrainment (Ecker et al., 2010; Hatori et al., 2008). However, what is the physiological function of ipRGC innervation in the SON region and what is the targeted neuron in the SON remain unclear.

Here, we demonstrate a visual circuit connecting the retina and the SON to modulate socio-sexual recognition memory (SSRM). Light exposure could activate GABAergic interneurons in the perinuclear zone of the SON (pSON), which in turn inhibit the oxytocin neurons in the SON and socio-sexual recognition memory. Genetically elimination of intrinsically photosensitive retinal ganglion cells (ipRGCs), or genetically silencing GABAergic interneurons in the pSON could block the light-induced SSRM reduction. Together, our results illustrate a direct sensory circuit that transmits the visual signal to modulate the oxytocin system in the brain and to regulate the formation of social memory.

## Material and Methods

### Animals

Adult C57BL/6J wild type (WT) mice were purchased from the National Laboratory Animal Center (NLAC), Taipei, Taiwan. Other transgenic mice used in this study were kept and bred at the Animal Facility of the Department of Life Science at National Taiwan University, Taipei, Taiwan. All transgenic mouse lines were maintained on a C57BL/6J background. Animals were housed under 12:12h light-dark cycle (lights on 0800-2000) at a temperature of 22°C, and had ad libitum access to normal chow and water unless otherwise stated. Experiments were approved by the Institutional Animal Ethics Committee of National Taiwan University, Taipei, Taiwan, and conducted in accordance with guidelines of the National Laboratory Animal Center. The RRID for Opn4Cre mouse is IMSR_JAX:035925, for Opn4DTA mouse is IMSR_JAX:035927, for Opn4CreERT2 mouse is IMSR_JAX:035926, for OyxotcinCre mouse is IMSR_JAX:024234, for ChR2-EYFP mouse is IMSR_JAX:024109, and for GAD-GFP mouse is IMSR_JAX:003718.

### Immunohistochemistry staining

Mice were deeply anesthetized with 1 mL (overdose) of Avertin (2, 2, 2-Tribromoethanol; Sigma Aldrich, 20 mg/mL) and perfused intracardially with 12 mL of iced PBS, following 25 mL of 4% paraformaldehyde (PFA). Brains were collected from the perfused animals and post-fixed in the same fixative at 4°C overnight. 80-μm-thick coronal brain sections were obtained using a vibratome (Campden Instruments, UK). For immunostaining, brain slices were incubated in blocking solution (6% goat serum in 1XPBST0.2%) for 2 hours at room temperature. Rabbit anti-oxytocin (Immunostar #1607001, 1: 4000) and Mouse IgG1α anti-cfos (AbCam ab208942, 1:1000) were diluted in blocking solution. Samples were incubated on an orbital shaker with primary antibody at 4°C overnight. Brain sections were washed and incubated in secondary antibody solution with Goat anti-Rabbit Alexa 568 (Biotium, 1:500) and Goat anti-Mouse IgG1α Alexa 488 (Invitrogen, 1:500) for 2 hours at room temperature. After washing, brain slices were mounted in mounting solution (Vectashield hard set with DAPI). Images were acquired using an inverted phase contrast fluorescence microscope (Zeiss Axio Observer Z1).

### Stereotaxic injection

Equipment used for stereotaxic surgery was properly sterilized. Animals were first anesthetized in gaseous isoflurane. The position of the head was secured by stereotaxic ear bars and tooth bars. Skull fur was shaved and skin was cleaned with iodine tincture and 70% ethanol. Incision was made to reveal the skull. After defining bregma and lambda, a small hole was drilled at regions of interest. The stereotaxic coordinates for the SON was -0.82 mm from bregma, ±1.35 mm lateral from midline, and 5.4 mm below the surface of the skull. For viral injection, glass needles with an inner diameter of 20 μm at the tip was filled with oil, and the needle was connected to a Hamilton syringe (#700, 5μL). 300 nL of virus was infused into the SON at a rate of 50 nL/min. GCaMP AAV vectors were obtained from addgene (Addgene_104492, and Addgene_100842) and packed in-house. After injection, the wound was stitched and animals were kept under warm light for recovery, with physical conditions monitored and body weight recorded for at least 3 days. For optic fiber implantation, the fiber was lowered into the brain at a speed of about 0.5 mm/min with a cannula holder (RWD, 68214, China). Then, dental cement (Hygenic, USA) was applied to secure the optic fiber to the skull. For unilateral implantation of optic fiber (for fiber photometry recordings), the stereotaxic coordinates were the same as the viral injection. For bilateral implantation (optogenetics), the fibers were tilted at a 10∘angle towards the midline. The stereotaxic coordinates for bilateral implantation at the SON was -0.82 mm from bregma, ±1.61 mm lateral from midline, and 5.25 mm below cortical surface.

### Behavior tests

Two trial social recognition tests followed the previous protocol. In brief, female mice were wild-type ordered from NLAC. Each female mouse was used for only once and only for a single male subject. All male subjects were single housed, kept under social isolation for at least 1 week. Before the experiment, the subject male and the stimuli female mice were kept under constant darkness (DD) for one day. On the experiment day, the two-trial social recognition test was conducted at CT12. During the test trial, the male subject was transferred to a novel cage and presented with a female for 10 minutes under dim light (5 lux). After the first trial, the male and female were transferred back to their home cage and kept under complete darkness (0 lux) for 1 hour. Then, the second trial was conducted in the same cage as the first trial with the same female stimulus. An hour of bright light treatment (800 lux) was given either before the social recognition test (CT11) or in between the two trials (CT12). During the test trials, the cage was recorded using an infrared camera.

### Fiber photometry recording

The fiber photometry recording system was custom-built based on the schematics on Thorlabs’ website (Thorlab, USA). The system consisted of 2 excitation sources, 405 nm and 470 nm LEDs. The LEDs were alternately turned on and off at 200 Hz in a square pulse pattern, which was driven by a PowerLab (ADInstruments, New Zealand). The illumination power of the LEDs was tuned to 0.4–0.9 mW/mm2 for the 405 nm LED and 0.7–1.4 mW/mm2 for the 470 nm LED at the tip of the optic fiber by LED drivers (Patel et al., 2020). Finally, GCaMP fluorescence was detected by a photodiode and the PowerLab at 1 kHz sampling rate. The fluorescence signal was processed by custom-written python codes referencing the open-source toolbox GuPPy (Sherathiya et al., 2021). During data processing, the signals were down-sampled to 100 Hz. The 405 nm LED exciting signal was used as the isobestic control. After both signals were low-pass filtered at 4 Hz, the isobestic control was fitted to the data (F470) using a least squares polynomial fit of degree 1, creating a fitted control (F_fitted control) as a result. ΔF/F was calculated by subtracting the fitted control from the data, then dividing by the fitted channel.

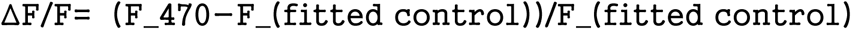

The subjects were kept under DD for 1 day until the recording. After tethering the optic fiber on the mice to the fiber photometry recording system, the animal was placed back to its cage and rested for 10 minutes with the LED excitation on. The whole process was completed under dim light (5 lux). The recording starts with 1 minute of baseline. In the second minute, a stimulus female was introduced to the cage to interact with the subject for 1 minute. Then, 2 dim-to-bright light transitions were recorded. In each transition, the two mice first interacted under dim light for 30 seconds, immediately followed by whole-cage illumination (800 lux) by white LED light for 30 seconds. Social interaction of the mice was video-recorded by an infrared camera.

### Optogenetic activation

Preparations for the social recognition test followed the description above. The male subject was tethered to the optic fiber cable at CT11 under darkness (0 lux) or bright light (800 lux) according to description. At CT11.5 to CT12, the subject was optogenetically stimulated with blue light (470 nm, 10 Hz, 10 ms duration). The light intensity was adjusted to 4 mW at the fiber tip. From CT12, the social recognition test proceeded as previously described.

## Results

### Light negatively influences socio-sexual recognition memory of male mice

To test whether light could influence social recognition memory in mice, we performed a two-trial social recognition test under dim light (5 lux) with a single stimulus female. Prior to the trials, wild-type male mice were subjected to an hour of light treatment (800 lux) or total darkness, designated as the L and D group, respectively (Figure 1A). We found that the recognition index of the L group was significantly lower than the D group, suggesting that light treatment had a negative effect on social recognition memory (Figure 1B-D, Extended Figure 1A and 1B). This phenomenon was specific to the opposite sex, as the recognition index of male-to-male interaction was not different in the L and D group (Extended Figure 1C and 1D). Furthermore, there was no significant difference between L and D group in the novel object recognition test (Extended Figure 2). Therefore, here we define this specific light-modulated recognition memory as socio-sexual recognition memory (SSRM). To test whether light could inadvertently alter the recognition index by inhibiting sociability during trial 1, we performed three-chamber sociability and social novelty tests for the L and D group. Neither sociability nor social novelty index was affected by an hour of light treatment (Extended Figure 3). Moreover, the investigation duration of trial 1 was not significantly different between the L and D group (Figure 1C, left panel). These results suggest that impaired SSRM in the L group was not due to a lack of socio-sexual interaction during trial 1. Next, we shifted the one-hour light treatment to the inter-trial interval to test whether light exposure could modulate SSRM by impairing memory retrieval (Figure 1D). If light exposure could modulate SSRM through memory retrieval but not memory formation, the recognition index should be lower in the inter-trial light exposure group compared to the D group. Our data showed that the recognition index was similar between inter-trial L and D groups (Figure 1E and 1F), suggesting that the SSRM retrieval is not affected by light exposure. Together, our data indicate that bright light exposure could modulate SSRM, potentially by impairing the memory formation stage.

**Figure 1.**
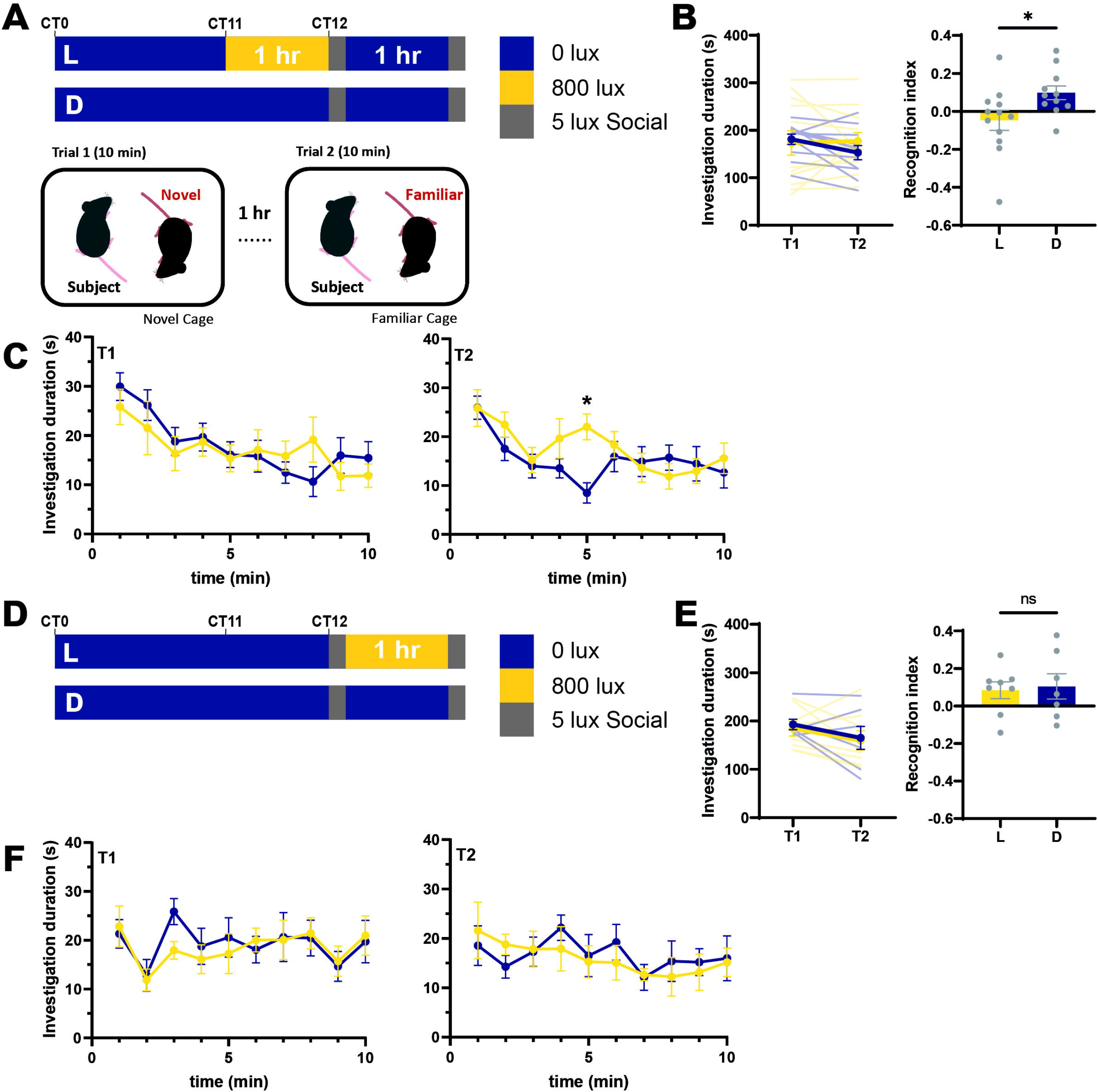
Pre-social light exposure impairs socio-sexual recognition memory (SSRM) (A) Schematics of pre-social light exposure and the two-trial social recognition test. (B) The investigation duration and recognition index of the light (L) and dark control (D) group. There is no significant difference in investigation duration at the first trial. However, there is a significant reduction in the recognition index in the L group compared to the D group. * indicates p = 0.0156, Mann-Whitney test. (C) The minute-wise investigation duration of L and D group. There is a significant increased investigation time only in L group 2^nd^ trial. * indicates p = 0.0106, Sidak’s multiple comparison test. (D) Inter-trial light exposure and social recognition test scheme. (E) There was no statistical significance between the L and D group in the investigation duration and recognition index. p = 0.9551, Mann-Whitney test. (F) The investigation duration per min from L and D groups.

**Figure 2.**
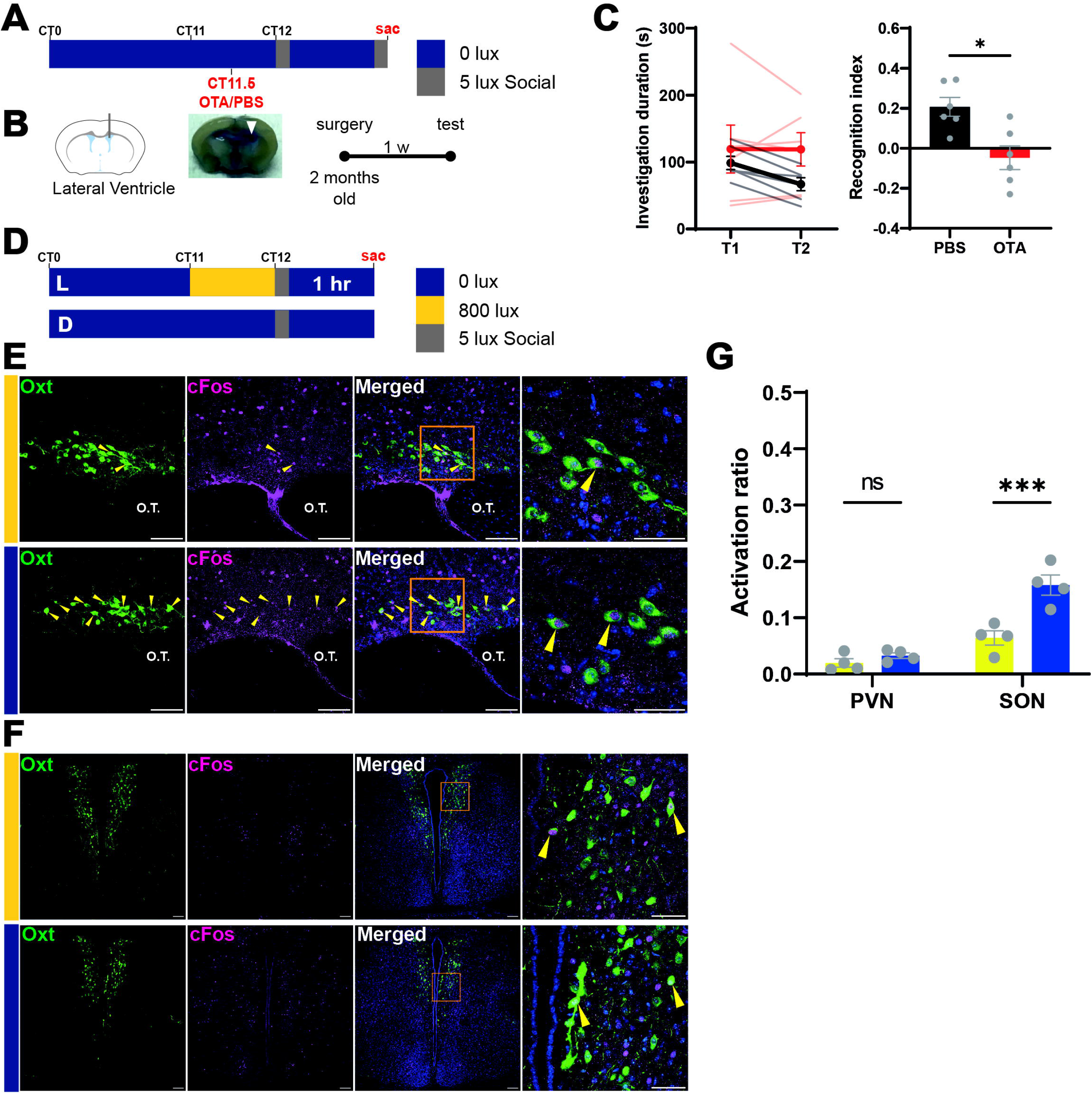
Supraoptic oxytocin neurons are inhibited by environmental luminance. **(A)** Schematics of social recognition test for the PBS-control and the OTA-treated group. **(B)** Representative photograph of brain slice with cannula implantation into the lateral ventricle, and dye verification of implantation site. **(C)** OTA significantly reduced the recognition index in mice compared to PBS injection control mice. * indicates p = 0.0260, Mann-Whitney test. **(D)** Schematic drawing of the c-fos staining experiment. **(E-F)** Representative confocal images of c-fos staining of oxytocin neurons in the SON (E) and PVN (F) in L and D group. Yellow arrows indicated oxytocin neurons co-labeled with cFos. For SON, high magnification images are 10 um Z-stack, low magnification images are 30 um Z-stack. Scale bar = 100 um for low magnification images and 50 um for high magnification images. **(G)** A significant reduction of c-Fos positive ratio in SON oxytocin neurons but not in PVN oxytocin neurons in the L group compared to the D group. *** indicates p = 0.0002, two-way ANOVA.

**Figure 3.**
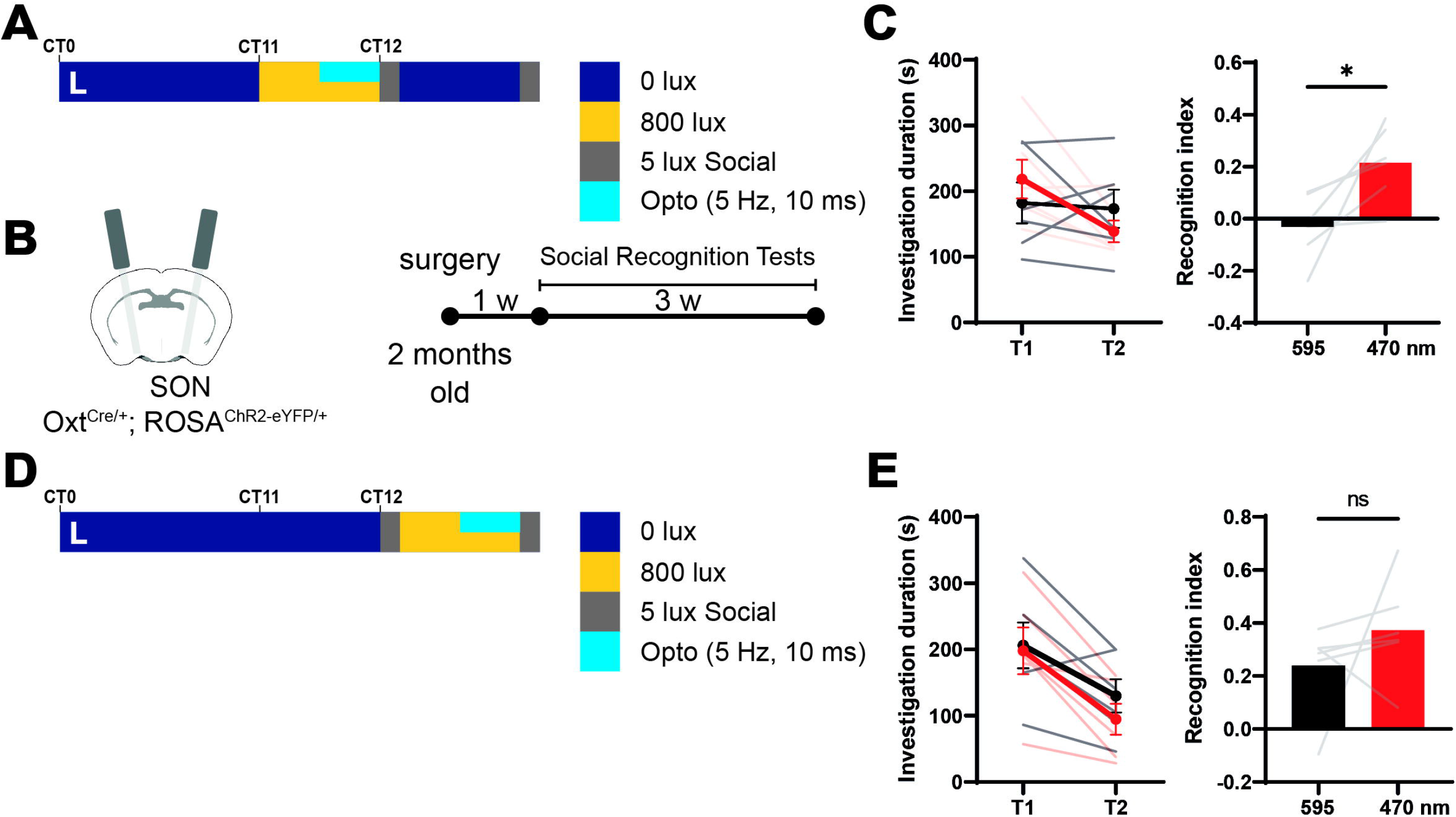
Optogenetic activation of SON^OT^ neurons under pre-social light treatment rescues SSRM. **(A-B)** Schematics of optogenetic activation of SON^OT^ neurons during pre-social light treatment. **(C)** The investigation duration and recognition index of the 595nm-control and the 470nm-activation group. There is a significant increase in recognition index in optogenetic activation group compared to control. * indicates p = 0.0313, Wilcoxon matched-pairs signed rank test. **(D)** Schematics of optogenetic activation of SON^OT^ neurons during inter-trial light treatment. **(E)** The investigation duration and recognition index of the 595nm-control and the 470nm-activation group. Optogenetic activation did not significantly change the recognition index. p = 0.1320, Wilcoxon matched-pairs signed rank test.

### Light inhibits SON oxytocin neurons to reduce SSRM

Oxytocin is a neurohormone involved in social interaction, pair bonding, and social memory formation. To confirm whether oxytocin is involved in SSRM in darkness, we performed an intracerebroventricular injection of the oxytocin receptor antagonist (OTA) through an implanted cannula 30 minutes before the test (Figure 2A and 2B). The recognition index of OTA-injected mice was significantly lower than PBS-injected control (Figure 2C). This result suggested that oxytocin signaling was involved in SSRM. In the hypothalamus, two main brain regions containing oxytocin neurons are the paraventricular nucleus (PVN) and supraoptic nucleus (SON). Anatomically, both PVN and SON are secondary targets of intrinsically photosensitive retinal ganglion cells (ipRGCs) through the suprachiasmatic nucleus (SCN) and peri-supraoptic nucleus (pSON), respectively. To investigate whether light could modulate oxytocin neural activation in the PVN and SON, we collected brain tissue from the mice 1 hour after socio-sexual interaction and performed c-Fos immunostaining from the light exposure and dark control groups (Figure 2D). We observed that the number of the c-fos positive SON^OT^ neurons from the L group is significantly lower compared to the D group (Figure 2E and 2G). On the other hand, there was no difference in the number of cFos positive PVN^OT^ neurons between the L and D group (Figure 2F and 2G). These results suggested that light may suppress the activity of SON^OT^ neurons but not PVN^OT^ neurons.

To test whether activation of SON^OT^ neurons could rescue the light-induced SSRM impairment, we implanted optic fiber above the SON in Oxt^Cre/+^; ROSA^ChR2-eYFP/+^ mice to artificially manipulate the activity of SON^OT^ neurons. Male mice with 1 hour of light exposure before the socio-sexual recognition test were performed similarly to the previous experimental setup. However, SON^OT^ neurons were exogenously activated by the 470 nm LED (5 Hz, 10 ms per pulse) during the second half of the light exposure period before the socio-sexual recognition test (Figure 3A and 3B). For negative control, 595 nm LED optostimulation (5 Hz, 10 ms per pulse) before the socio-sexual recognition test was performed on a separate experiment day. The recognition index for the 470 nm opto-stimulation group was significantly higher than the 595 nm control group (Figure 3C), suggesting that optogenetic activation of SON^OT^ neurons can rescue light-induced reduction of SSRM. To investigate whether activation of SON^OT^ neurons could modulate SSRM through memory formation or retrieval, we optogenetically activated SON^OT^ neurons during the second half of the inter-trial light treatment (Figure 3D). Similar to the previous result, the recognition index was not different between the 595 nm control and the 470 nm-stimulation groups (Figure 3E). These results indicate that the activation of SON^OT^ neurons could alleviate light induced SSRM reduction by potentially modulating the memory formation process.

### Brn3b^+^ M1 ipRGCs transmit environmental light information to the SON

It has been shown that distinct subtypes of ipRGC could project to different brain regions to modulate various non-image forming functions. Specifically, Brn3b negative M1 ipRGCs project to the SCN, whereas Brn3b positive M1 and non-M1 ipRGCs project to other brain regions. Therefore, we next want to test whether ipRGCs and which sub-populations of ipRGCs are involved in the light-induced suppression of SON^OT^ neurons. First, we performed c-Fos staining of SON^OT^ neurons using the same experimental condition as previously described but in Opn4^DTA/DTA^ mice, whose M1 ipRGCs were genetically ablated. We found that c-Fos positive SON^OT^ neurons were similar between the L and D groups (Extended Figure 4A-C), suggesting that M1 ipRGCs are required for light-dependent modulation of SON^OT^ neurons. To confirm ipRGCs could modulate SON^OT^ neurons in a direct circuit but not from the putative SCN-PVN-SON circuit, we performed c-Fos staining in Opn4^Cre/+^; Brn3b^zDTA/+^ mice. In this mouse line, the ipRGC to SCN circuit and circadian photoentrainment remains functional while the ipRGC to other brain areas such as the pSON are eliminated. We found that the ratio of c-Fos positive SON^OT^ neurons was also similar between the L and D group in Opn4^Cre/+^; Brn3b^zDTA/+^ mice (Extended Figure 4D-F). Together, these results indicate that SON-projecting Brn3b positive M1 ipRGCs are the primary retinal input that modulates the activity of the SON^OT^ neurons. Finally, to confirm the functional connection between ipRGCs and SON^OT^ neurons *in vivo*, we injected AAV9/flox-GCaMP6s at the SON region and implanted optic fiber above the SON in Oxt^Cre/+^ and Oxt^Cre/+^; Opn4^DTA/DTA^ mice to record the calcium response of SON^OT^ neurons using fiber photometry (Figure 4A and 4B). As expected, the GCaMP signal from SON^OT^ neurons was suppressed by bright (800 lux) light exposure compared to the baseline in Oxt^Cre/+^ mice. The GCaMP signal during the 30 sec of light exposure was lower than the baseline period (Figure 4C, black trace). Furthermore, the GCaMP signal remained unchanged during the light exposure period in the M1 ipRGC eliminated Opn4^DTA/DTA^ mice (Figure 4C, red trace). The difference of mean GCaMP signal in control mice between baseline and light exposure was significantly larger than the difference in Opn4^DTA/DTA^ mice, which was close to zero (Figure 4D). Together, our results indicate that Brn3b positive M1 ipRGCs are necessary for transmitting light information to suppress the activity of SON^OT^ neurons acutely.

**Figure 4.**
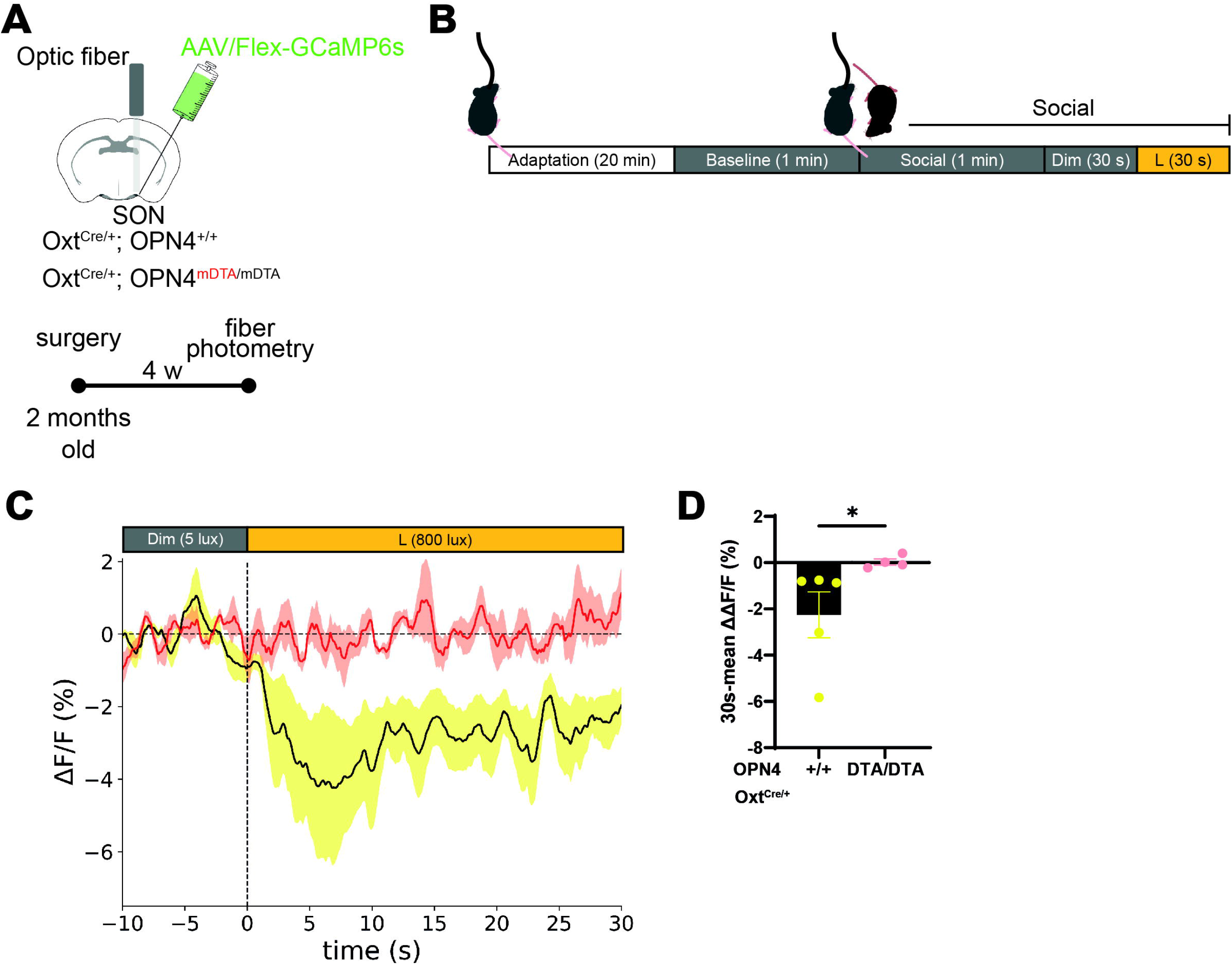
SON^OT^ neuronal activity is suppressed by environmental luminance. **(A-B)** Schematics of fiber photometry recording. **(C)** Average delta-F/F signal trace from SON oxytocin neurons with 10 sec of dim light (5 lux) followed by 30 sec of bright light (800 lux) in OPN4^DTA/DTA^; Oxt^Cre/+^ mice (red line) and OPN4^+/+^; Oxt^Cre/+^ mice (black line). The color shade indicates standard derivation. **(D)** Mean change of delta-F/F in (C). There is a significant reduction of GCaMP signal in control mice after light exposure. (*p = 0.0159, Mann-Whitney test).

### SON-projecting ipRGCs mediate modulation of socio-sexual recognition by light

Next, we wanted to ask whether melanopsin signaling and Brn3b positive M1 ipRGCs are required for light-induced modulation of SSRM behaviorally (Figure 5A). In control (Opn4^Cre/+^) mice, the recognition index of the L group was significantly lower than the D group, similar to WT mice (Figure 5B). In contrast, the recognition indexes were similar between the L and D group in Opn4^DTA/DTA^ and Opn4^Cre/+^; Brn3b^zDTA/+^ animals (Figure 5C and 5D). These results suggest that Brn3b positive M1 ipRGCs, but not SCN-projecting M1 ipRGC, are required for light-induced reduction of the SSRM. Strikingly, in melanopsin knockout (MKO) animals, the recognition index did not significantly differ between the L and D groups (Figure 5E). It suggests that the melanopsin photo-detection system may be required for prolonged inhibition of SON^OT^ neurons and the subsequent light-induced SSRM reduction. To confirm that signal from SON targeting ipRGCs are sufficient to modulate SSRM *in vivo*, we express channelrhodopsin in ipRGCs using Opn4^Cre-ERT2/+^; ROSA^ChR2-eYFP/+^ mice and activate their terminal at the pSON with 470 nm LED light stimulation (10 Hz, 10 ms) through the optic fiber implanted above the SON (Figure 5F and 5G). The recognition index of the 470 nm-stimulated trials was significantly lower than the control 595 nm optostimulation trials (Figure 5H), indicating that activation of SON-targeting ipRGCs is sufficient to reduce SSRM. Together, our results suggest that ipRGCs projecting to the pSON are necessary and sufficient to modulate the SSRM by inhibiting the SON^OT^ neurons.

**Figure 5.**
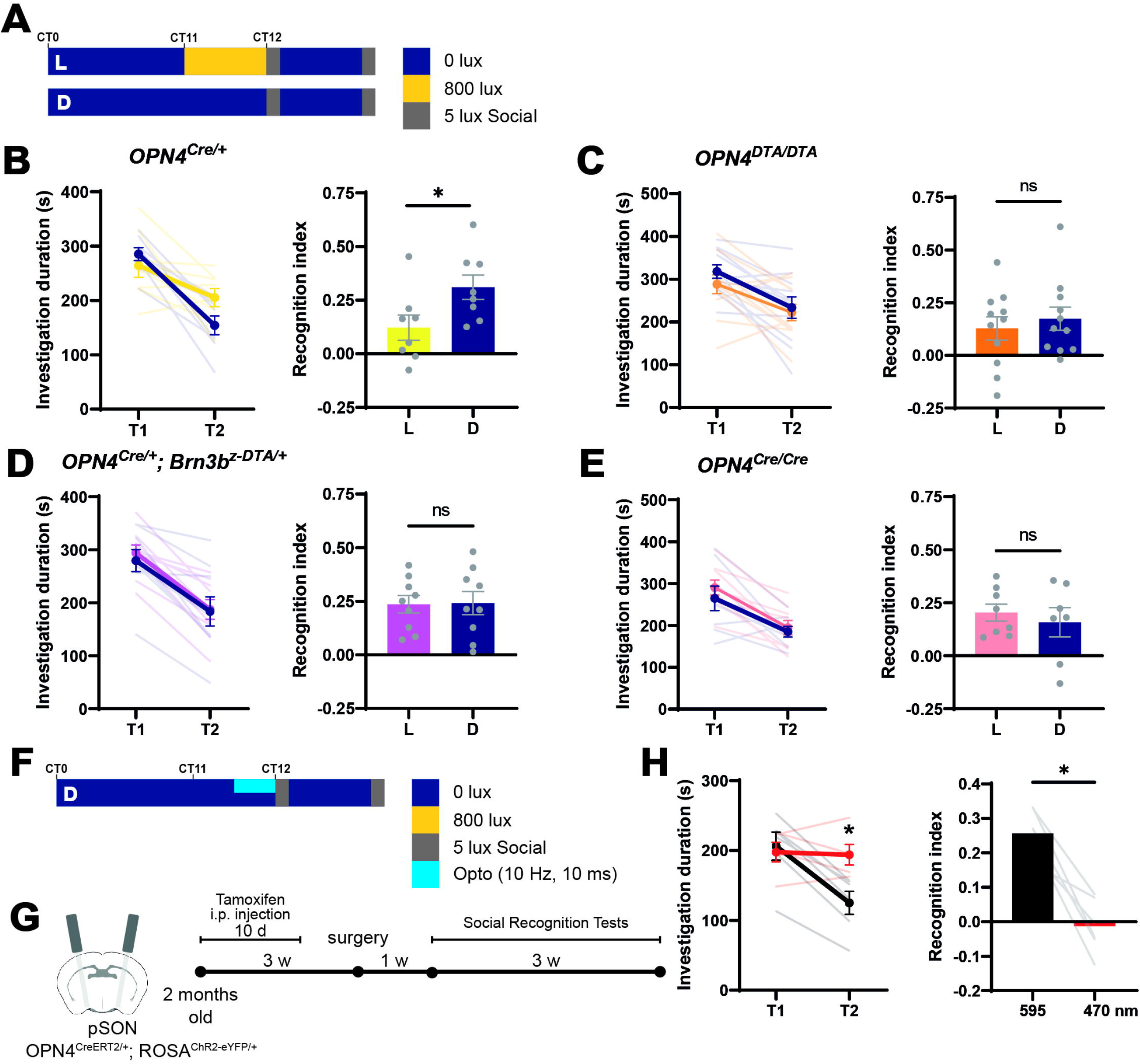
SON-projecting ipRGCs are required and sufficient to drive light-induced impairment of SSRM. **(A)** Schematics of pre-social light exposure and two-trial social recognition test for each genotype. **(B)** Light exposure significantly reduces the recognition index in control Opn4^Cre/+^ mice, similar to WT mice. (*p = 0.0499, Mann-Whitney test). **(C-E)** There is no significant different in recognition index between light exposure and dark control groups in OPN4^DTA/DTA^ (C, p = 0.7969), OPN4^Cre/+^; Brn3b^z-DTA/+^ (D, p > 0.9999), and OPN4^Cre/Cre^ mice (E, p = 0.9551) using Mann-Whitney test. **(F-G)** Schematics of optogenetic activation of SON-projecting ipRGCs under darkness. **(H)** Optogenetic activation of ipRGCs terminal at SON with 470 nm light significantly enhances the recognition index compared to 595 nm control. * indicates p = 0.0313, Wilcoxon matched-pairs signed rank test.

### ipRGCs synapse on GABAergic neurons in the perinuclear zone of the SON

Since previous literature suggested that SON^OT^ neurons could be inhibited by GABAergic interneurons (Brussaard et al., 1997; Engelmann et al., 2004; Roland & Sawchenko, 1993; Theodosis et al., 1986), it is possible that ipRGCs may indirectly inhibit SON^OT^ neurons through GABAergic neurons in the pSON. To examine the downstream connection of ipRGCs in the SON/pSON region, we performed triple labeling using GAD67^eGFP/+^ to label GABAergic neurons, CTB-Alexa 568 injection in the eye to label ipRGC terminals in the pSON, and immunostaining of synaptophysin to label the presynaptic site (Extended Figure 5A). We observed that ipRGC axon terminals are mainly located at the peri-SON and form many putative synaptic contacts on the soma of GABAergic neurons (Extended Figure 5B-F). This data suggests that ipRGCs may indirectly suppress SON^OT^ neurons through GABAergic interneurons in the peri-SON region. To test whether light exposure could activate GABAergic neuron near SON *in vivo*, we again perform GCaMP recording in vGAT^Cre/+^ mice injected with AAV9/flox-GCaMP7f using fiber-photometry (Figure 6A and 6B). Light exposure significantly increases GCaMP signal compared to dark baseline (Figure 6C and 6D). Next, to confirm that GABAergic neurons in the pSON could modulate SSRM, we selectively activated GABAergic neurons optogenetically through an implanted optic fiber above the pSON in vGAT^Cre/+^; ROSA^ChR2-eYFP/+^ mice (Figure 6E and 6F). The recognition index of the 470 nm-stimulated trails was significantly lower than the control 595 nm stimulated trails (Figure 6G). Furthermore, elimination of neurotransmitter release from these GABAergic neurons using vGAT^Cre/+^ mice injected with AAV9-Flex-TeLC in the SON could also alleviate the light-induced SSRM reduction (Figure 6H-J). Together, our results indicate that light could activate pSON GABAergic neurons to modulate SSRM.

**Figure 6.**
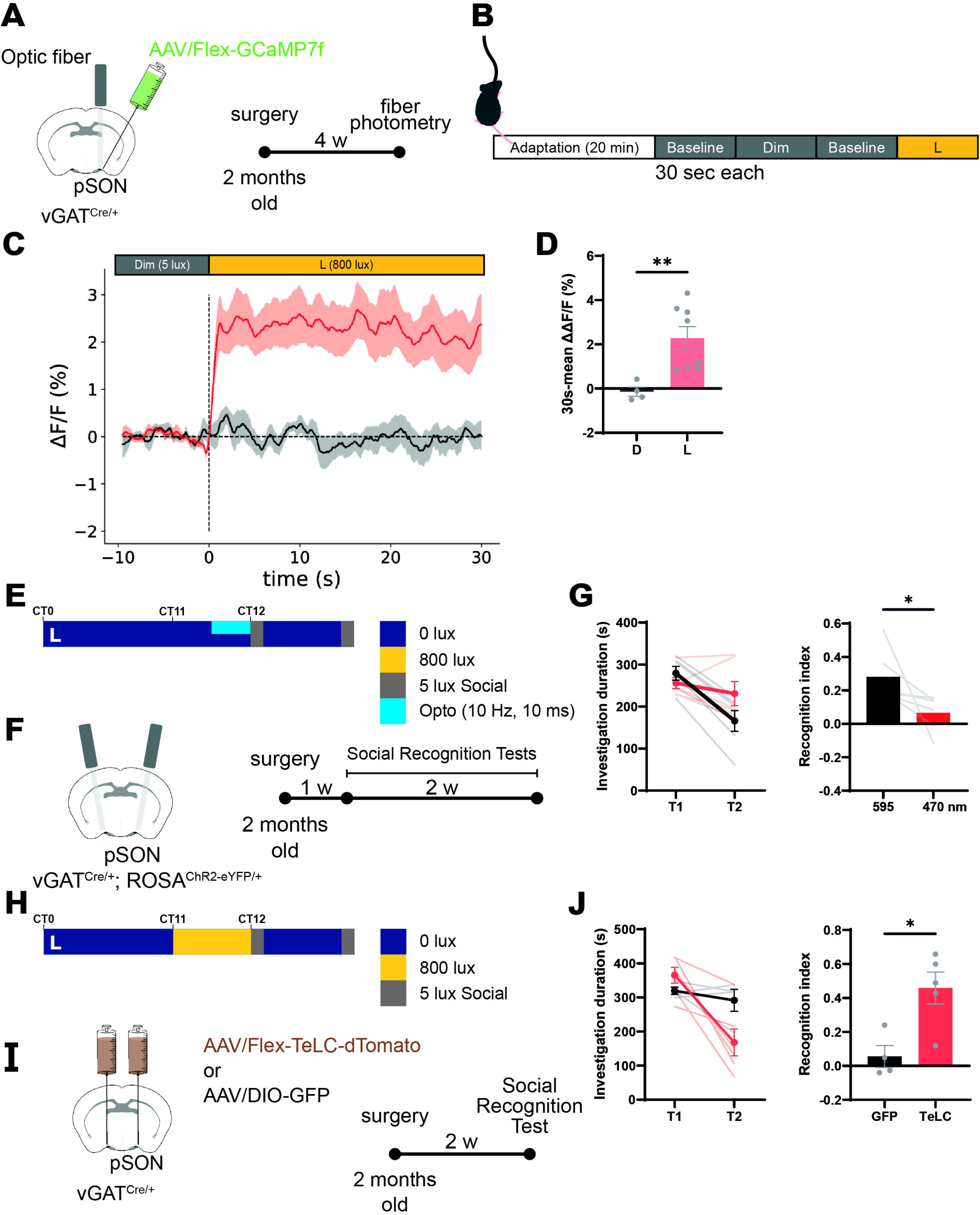
GABAergic neurons in the perinuclear zone of the SON are required and sufficient to drive light-induced impairment of SSRM. **(A-B)** Schematics of fiber photometry recording. **(C)** Average delta-F/F signal trace from pSON GABAergic neurons during the transition from baseline to dim light (5 lux, black line) and from baseline to bright light (800 lux, red line) in vGAT^Cre/+^ mice. The color shade indicates standard derivation. **(D)** Mean change of delta-F/F between transition from (C). Light exposure significantly increased the delta-F/F in GABAergic neurons at pSON region. * indicates p = 0.0062, unpaired t test. **(E-F)** Schematics of optogenetic activation of GABAergic neurons in the pSON in darkness. **(G)** The investigation duration and recognition index of the 595nm-control and the 470nm-activation group. Optogenetic activation of GABAergic neurons at pSON region significantly decreased the recognition index. * indicates p = 0.0313, Wilcoxon matched-pairs signed rank test. **(H-I)** Schematics of silencing GABAergic neurons in the pSON and social recognition test with light treatment. **(J)** The investigation duration and recognition index of the GFP-control and the TeLC-silencing group. Mice with genetic silencing GABAergic neurons at pSON region had higher recognition index than control mice. * indicates p = 0.0317, Mann-Whitney test).

## Discussion

Previous studies have shown that light can downregulate object and odor recognition memory performance in an intensity-dependent manner (Hasan et al., 2021). However, whether light can modulate other types of memory is unknown. Here we show that optogenetic activation of ipRGC terminals or GABAergic neurons in the pSON could inhibit SSRM. In contrast, optogenetic activation of oxytocin neurons in the SON could increase SSRM. Therefore, it is likely that ipRGC may activate GABAergic interneurons at the pSON, which inhibit SON^OT^ neurons. Therefore, our results propose a novel neuronal circuity that conveys the light signal to inhibit SSRM through the ipRGC-pSON-SON^OT^ pathway. This circuit is necessary and sufficient to transmit external light information to influence the oxytocin system in the brain, which provides a functional input from the visual system to modulate oxytocin neurons and social memory directly through the RHT pathway.

With pre-social bright light treatment, naïve wild-type male mice showed a significantly lower recognition index than dark control mice. A reduced recognition index could theoretically originate from acute inhibition of social interaction, impairment in memory formation, or impairment of memory recall. Acute inhibition of social interaction in the first trial may influence the initial stage of social recognition memory formation. However, additional sociability and social novelty tests both show similar indexes between the L and D groups. Furthermore, we did not observe a significant difference in the interaction duration between the L and D groups in the first trial. Therefore, our data suggests that the reduction of the recognition index after light exposure is unlikely due to an acute inhibition of socio-sexual investigation duration. Next, since light exposure and optogenetic activation of SON^OT^ neurons could only affect the recognition index before the test but not during the inter-trail interval, these results indicate that light stimulation may primarily modulate the formation of SSRM but not inhibit the recall of SSRM. In line with this idea, Ferguson et al. reported that oxytocin injection in oxytocin-knockout animals before initial social exposure could greatly promote social memory, whereas injection after social exposure did not have any effect on social memory (Ferguson et al., 2001). They speculated that the presence of oxytocin during the acquisition phase assists in the olfactory processing of social cues, but its presence is not required for recalling social memory. Together our study strongly suggests that light might inhibit SON^OT^ neurons and further reduce the acquisition phase of SSRM. A recent study showed that ipRGC could activate SON^OT^ neurons at the early postnatal stage (Hu et al., 2022). Although they showed the opposite effect of light on oxytocin content in the brain than this study, our experiments were carried out at the adult stage after sexual maturation. Therefore, there are two potential hypotheses to explain these contradicted results. First, the GABAergic circuits may serve as a positive stimulation for oxytocin neurons in the SON during the early postnatal day. Therefore, light could activate SON^OT^ neurons through the same pathway proposed by this study at the early postnatal stage. Second, the neural circuits from ipRGCs to the SON^OT^ neurons are remodeled after the early postnatal stage. Further study to show the change of ipRGC circuity functionally or anatomically may provide additional insight into the switch.

Previously, Lister and Hilakivi observed that in an unfamiliar arena, high luminance suppressed social interaction between male Swiss mice (Lister & Hilakivi, 1988). However, here we specifically measure the male to female investigation time. Therefore, the interaction between the opposite sex may explain the difference between our result and previous literature. Alternatively, habituation duration in the testing chamber could also influence the investigation time. In our study, male mice only spend 50-100 sec investigation time on another male mouse compared to 200-300 sec on female mice. The investigation time during the first trial period may not be sufficient to form a robust social recognition memory under our experimental setup. Together, our study primarily indicates that light could strongly influence the socio-sexual recognition memory. Whether social recognition memory between a pair of male mice could be influenced by light may require further study using a different testing protocol.

It has been shown that oxytocin is an important signal for social recognition. Without oxytocin, male mice display impaired recognition to female mice (Ferguson et al., 2000). In this study, we showed that ipRGCs could suppress the activity of SON^OT^ neurons specifically after light exposure by c-fos staining and *in vivo* calcium imaging with fiber photometry. In support of our finding, Devarajan and Rusak reported that oxytocin concentration can be suppressed by a nocturnal light pulse (Devarajan & Rusak, 2004), which may reflect decreased oxytocin neural activity. In contrast to previous literature, which showed contact and sound inputs could activate PVN^OT^ neurons, we observed that light exposure affects only the activation ratio of SON^OT^ neurons, but not PVN^OT^ neurons. Interestingly, the neuronal input for PVN^OT^ regulation by touch or sound primarily originated from the somatosensory and auditory cortex respectively. However, here we showed that light could modulate SON oxytocin neuronal activity and SSRM. Since a specific pattern of physical contact and sound, such as soft touch and pup calling during parenting, could strongly promote oxytocin release, a special modality of such sensory input and cortical computation could be required to activate oxytocin neurons. For light, the activation of ipRGC with melanopsin photo-response may directly inhibit the oxytocin neurons in the SON, suggesting that a simple luminance signal is adequate to modulate the oxytocin system and their downstream physiological functions. Furthermore, the PVN and SON may serve as different target sites for distinct origins of signaling to modulate the oxytocin system.

It has been shown that ipRGC comprises many distinct subpopulations from M1-M6 according to their morphological and electrical properties. Furthermore, genetic markers such as Brn3b, glycine, and GABA could also separate ipRGCs into distinct functional groups. In contrast to wild-type animals, the activation ratio of SON^OT^ neurons according to cFos staining in Opn4^DTA/DTA^ as well as Opn4^Cre/+^; Brn3b^zDTA/+^ mice did not differ in the L and D group. Recognition indexes were also similar between the L and D group in two different ipRGC eliminated mice. Previously, Li and Schmidt observed that ipRGC innervation at the SON is absent in Opn4^Cre/tau-LacZ^; Brn3b^z-dta/+^ mice (Li & Schmidt, 2018). Together, our results suggest that Brn3b^+^ M1 ipRGCs are the primary retinal input to the SON to modulate the activity of oxytocin neurons. Without ipRGC projection, SON^OT^ neural activity and SSRM are no longer modulated by environmental light. Interestingly, light exposure could not modulate SSRM in the absence of melanopsin with ipRGCs intact. This indicates that melanopsin photo-detection is required for sustained inhibition of SON^OT^ neurons. This phenomenon differs from other ipRGC-dependent functions such as circadian photo-entrainment, in which rod and cone signals could compensate for the lack of melanopsin. Since the melanopsin photo-detection system is long-lasting and weakly adapted to continuous light exposure compared to rods and cones (Berson et al., 2002), prolonged activation of the ipRGC-SON circuit is likely required to modulate the oxytocin and SSRM. Together, our results indicate an important function of melanopsin photopigment in modulating our physiological function.

Although most ipRGCs release glutamate and PACAP as their neurotransmitter, a recent study showed that a subset of ipRGCs could co-release GABA in addition to glutamate in the SCN (Takuma Sonoda, 2020). However, another study showed that SON-innervating ipRGCs only release glutamate (Berry et al., 2022). Here we found that ipRGCs also inhibited SON^OT^ neurons after light exposure. It is possible that ipRGC may inhibit SON^OT^ neurons through direct contact with GABA release or through an indirect pathway. Our data showed that RGC terminals make many synaptic contacts on the soma of GABAergic neurons in the pSON. Various studies have shown that GABAergic input to the SON can decrease the firing rate of oxytocin neurons (Brussaard et al., 1997; Engelmann et al., 2004; Lee et al., 2015). In this study, our data did not completely eliminate a potential small parallel pathway in which ipRGCs directly inhibit SON^OT^ neurons through GABA release. However, we showed that activation of pSON^GABA^ neurons is sufficient and necessary to modulate SSRM behaviorally. We believe that the pSON GABAergic population may relay the ipRGC signal and convert it to inhibit SON^OT^ neurons, thereby impairing memory. Together, our study reveals a functional circuitry originating from ipRGC to modulate the oxytocin system in the SON. Therefore, in addition to touch and sound sensory inputs, luminance signals from the visual system could also modulate the SSRM and potentially other oxytocin-related physiological functions. Finally, our proposed ipRGC, pSON GABAergic neurons, and SON oxytocin neurons circuit may provide a parallel pathway in the visual system to modulate oxytocin content in the brain. Thus, the canonical vision input could function as the salient interaction stimulation during the social interaction to regulate social behaviors.

## Acknowledgments

This work was supported by the Taiwan National Science and Technology Council grant NTSC 106-2311-B-002-033-MY3 and 111-2636-B-002-021 (to S.-K.C.). We thank the Technology Commons, College of Life Science at National Taiwan University for technical assistance with confocal imaging. The authors declare no competing financial conflicts of interest.

## Author contributions

P-Y. L. and S-K. C. conceptualize the project. Y.-F. H., P-Y. L. and J.-H. Y. performed all experiments and analyses. Y.-F. H. and S-K. C. wrote the manuscript.

## Reference

Armstrong, W. E. (2015). Chapter 14 - Hypothalamic Supraoptic and Paraventricular Nuclei. In G. Paxinos (Ed.), The Rat Nervous System (Fourth Edition) (pp. 295–314). Academic Press. https://doi.org/10.1016/B978-0-12-374245-2.00014-0

Badia, P., Myers, B., Boecker, M., Culpepper, J., & Harsh, J. R. (1991). Bright light effects on body temperature, alertness, EEG and behavior. Physiology & Behavior, 50(3), 583–588. https://doi.org/10.1016/0031-9384(91)90549-4

Berry, M. H., Moldavan, M., Garrett, T., Meadows, M., Cravetchi, O., White, E., von Gersdorff, H., Wright, K. M., Allen, C., & Sivyer, B. (2022). A subtype of melanopsin ganglion cells encodes ground luminance. bioRxiv, 2022.2004.2006.485949. https://doi.org/10.1101/2022.04.06.485949

Berson, D. M., Dunn, F. A., & Takao, M. (2002). Phototransduction by retinal ganglion cells that set the circadian clock. Science, 295(5557), 1070–1073. http://www.ncbi.nlm.nih.gov/entrez/query.fcgi?cmd=Retrieve&db=PubMed&dopt=Citation&list_uids=11834835

Brussaard, A. B., Kits, K. S., Baker, R. E., Willems, W. P. A., Leyting-Vermeulen, J. W., Voorn, P., Smit, A. B., Bicknell, R. J., & Herbison, A. E. (1997). Plasticity in Fast Synaptic Inhibition of Adult Oxytocin Neurons Caused by Switch in GABAA Receptor Subunit Expression. Neuron, 19(5), 1103–1114. https://doi.org/10.1016/S0896-6273(00)80401-8

Cajochen, C., Zeitzer, J. M., Czeisler, C. A., & Dijk, D.-J. (2000). Dose-response relationship for light intensity and ocular and electroencephalographic correlates of human alertness. Behavioural Brain Research, 115(1), 75–83.

Carcea, I., Caraballo, N. L., Marlin, B. J., Ooyama, R., Riceberg, J. S., Mendoza Navarro, J. M., Opendak, M., Diaz, V. E., Schuster, L., Alvarado Torres, M. I., Lethin, H., Ramos, D., Minder, J., Mendoza, S. L., Bair-Marshall, C. J., Samadjopoulos, G. H., Hidema, S., Falkner, A., Lin, D., … Froemke, R. C. (2021). Oxytocin neurons enable social transmission of maternal behaviour. Nature, 596(7873), 553–557. https://doi.org/10.1038/s41586-021-03814-7

Chellappa, S. L., Steiner, R., Oelhafen, P., Lang, D., Götz, T., Krebs, J., & Cajochen, C. (2013). Acute exposure to evening blue-enriched light impacts on human sleep. Journal of Sleep Research, 22(5), 573–580. https://doi.org/10.1111/jsr.12050

Chen, S. K., Badea, T. C., & Hattar, S. (2011). Photoentrainment and pupillary light reflex are mediated by distinct populations of ipRGCs. Nature, 476(7358), 92–95. https://doi.org/10.1038/nature10206

Devarajan, K., & Rusak, B. (2004). Oxytocin levels in the plasma and cerebrospinal fluid of male rats: effects of circadian phase, light and stress. Neuroscience Letters, 367(2), 144–147. https://doi.org/10.1016/j.neulet.2004.05.112

Ecker, J. L., Dumitrescu, O. N., Wong, K. Y., Alam, N. M., Chen, S. K., Legates, T., Renna, J. M., Prusky, G. T., Berson, D. M., & Hattar, S. (2010). Melanopsin-Expressing Retinal Ganglion-Cell Photoreceptors: Cellular Diversity and Role in Pattern Vision. Neuron, 67(1), 49–60. https://doi.org/S0896-6273(10)00419-8 [pii] 10.1016/j.neuron.2010.05.023

Eliava, M., Melchior, M., Knobloch-Bollmann, H. S., Wahis, J., da Silva Gouveia, M., Tang, Y., Ciobanu, A. C., Triana Del Rio, R., Roth, L. C., Althammer, F., Chavant, V., Goumon, Y., Gruber, T., Petit-Demouliere, N., Busnelli, M., Chini, B., Tan, L. L., Mitre, M., Froemke, R. C., … Grinevich, V. (2016). A New Population of Parvocellular Oxytocin Neurons Controlling Magnocellular Neuron Activity and Inflammatory Pain Processing. Neuron, 89(6), 1291–1304. https://doi.org/10.1016/j.neuron.2016.01.041

Engelmann, M., Bull, P. M., Brown, C. H., Landgraf, R., Horn, T. F. W., Singewald, N., Ludwig, M., & Wotjak, C. T. (2004). GABA selectively controls the secretory activity of oxytocin neurons in the rat supraoptic nucleus. European Journal of Neuroscience, 19(3), 601–608. https://doi.org/10.1111/j.1460-9568.2004.03151.x

Ferguson, J. N., Aldag, J. M., Insel, T. R., & Young, L. J. (2001). Oxytocin in the Medial Amygdala is Essential for Social Recognition in the Mouse. The Journal of Neuroscience, 21(20), 8278. https://doi.org/10.1523/JNEUROSCI.21-20-08278.2001

Ferguson, J. N., Young, L. J., Hearn, E. F., Matzuk, M. M., Insel, T. R., & Winslow, J. T. (2000). Social amnesia in mice lacking the oxytocin gene. Nature Genetics, 25(3), 284–288. https://doi.org/10.1038/77040

Fernandez, D. C., Fogerson, P. M., Lazzerini Ospri, L., Thomsen, M. B., Layne, R. M., Severin, D., Zhan, J., Singer, J. H., Kirkwood, A., Zhao, H., Berson, D. M., & Hattar, S. (2018). Light Affects Mood and Learning through Distinct Retina-Brain Pathways. Cell, 175(1), 71–84 e18. https://doi.org/10.1016/j.cell.2018.08.004

Hasan, S., Tam, S. K. E., Foster, R. G., Vyazovskiy, V. V., Bannerman, D. M., & Peirson, S. N. (2021). Modulation of recognition memory performance by light and its relationship with cortical EEG theta and gamma activities. Biochemical Pharmacology, 191, 114404. https://doi.org/10.1016/j.bcp.2020.114404

Hatori, M., Le, H., Vollmers, C., Keding, S. R., Tanaka, N., Buch, T., Waisman, A., Schmedt, C., Jegla, T., & Panda, S. (2008). Inducible ablation of melanopsin-expressing retinal ganglion cells reveals their central role in non-image forming visual responses. PLoS ONE, 3(6), e2451.

Hattar, S., Kumar, M., Park, A., Tong, P., Tung, J., Yau, K.-W., & Berson, D. M. (2006). Central projections of melanopsin-expressing retinal ganglion cells in the mouse. Journal of Comparative Neurology, 497(3), 326–349. https://doi.org/10.1002/cne.20970

Hu, J., Shi, Y., Zhang, J., Huang, X., Wang, Q., Zhao, H., Shen, J., Chen, Z., Song, W., Zheng, P., Zhan, S., Sun, Y., Cai, P., An, K., Ouyang, C., Zhao, B., Zhou, Q., Xu, L., Xiong, W., … Xue, T. (2022). Melanopsin retinal ganglion cells mediate light-promoted brain development. Cell, 185(17), 3124-3137.e3115. https://doi.org/10.1016/j.cell.2022.07.009

Hughes, R. N., Hancock, N. J., Henwood, G. A., & Rapley, S. A. (2014). Evidence for anxiolytic effects of acute caffeine on anxiety-related behavior in male and female rats tested with and without bright light. Behavioural Brain Research, 271, 7–15. https://doi.org/10.1016/j.bbr.2014.05.038

Lee, S. W., Kim, Y.-B., Kim, J. S., Kim, W. B., Kim, Y. S., Han, H. C., Colwell, C. S., Cho, Y.-W., & In Kim, Y. (2015). GABAergic inhibition is weakened or converted into excitation in the oxytocin and vasopressin neurons of the lactating rat. Molecular Brain, 8(1), 34. https://doi.org/10.1186/s13041-015-0123-0

LeGates, T. A., Altimus, C. M., Wang, H., Lee, H.-K., Yang, S., Zhao, H., Kirkwood, A., Weber, E. T., & Hattar, S. (2012). Aberrant light directly impairs mood and learning through melanopsin-expressing neurons. Nature, 491(7425), 594–598. https://doi.org/10.1038/nature11673

Li, J. Y., & Schmidt, T. M. (2018). Divergent projection patterns of M1 ipRGC subtypes. Journal of Comparative Neurology, 526(13), 2010–2018. https://doi.org/10.1002/cne.24469

Liao, P.-Y., Chiu, Y.-M., Yu, J.-H., & Chen, S.-K. (2020). Mapping Central Projection of Oxytocin Neurons in Unmated Mice Using Cre and Alkaline Phosphatase Reporter [Original Research]. Frontiers in Neuroanatomy, 14. https://doi.org/10.3389/fnana.2020.559402

Lister, R. G., & Hilakivi, L. A. (1988). The effects of novelty, isolation, light and ethanol on the social behavior of mice. Psychopharmacology, 96(2), 181–187. https://doi.org/10.1007/BF00177558

Ludwig. (1998). Dendritic Release of Vasopressin and Oxytocin. Journal of Neuroendocrinology, 10(12), 881–895. https://doi.org/10.1046/j.1365-2826.1998.00279.x

Lukas, M., Toth, I., Reber, S. O., Slattery, D. A., Veenema, A. H., & Neumann, I. D. (2011). The Neuropeptide Oxytocin Facilitates Pro-Social Behavior and Prevents Social Avoidance in Rats and Mice. Neuropsychopharmacology, 36(11), 2159–2168. https://doi.org/10.1038/npp.2011.95

Lupi, D., Oster, H., Thompson, S., & Foster, R. G. (2008). The acute light-induction of sleep is mediated by OPN4-based photoreception. Nature Neuroscience, 11(9), 1068–1073. https://doi.org/10.1038/nn.2179

Milosavljevic, N., Cehajic-Kapetanovic, J., Procyk, C. A., & Lucas, R. J. (2016). Chemogenetic Activation of Melanopsin Retinal Ganglion Cells Induces Signatures of Arousal and/or Anxiety in Mice. Current biology : CB, 26(17), 2358–2363. https://doi.org/10.1016/j.cub.2016.06.057

Nakajima, M., Görlich, A., & Heintz, N. (2014). Oxytocin Modulates Female Sociosexual Behavior through a Specific Class of Prefrontal Cortical Interneurons. Cell, 159(2), 295–305. https://doi.org/10.1016/j.cell.2014.09.020

Oettl, L.-L., Ravi, N., Schneider, M., Scheller Max F., Schneider, P., Mitre, M., da Silva Gouveia, M., Froemke Robert C., Chao Moses V., Young, W. S., Meyer-Lindenberg, A., Grinevich, V., Shusterman, R., & Kelsch, W. (2016). Oxytocin Enhances Social Recognition by Modulating Cortical Control of Early Olfactory Processing. Neuron, 90(3), 609–621. https://doi.org/10.1016/j.neuron.2016.03.033

Resendez, S. L., Namboodiri, V. M. K., Otis, J. M., Eckman, L. E. H., Rodriguez-Romaguera, J., Ung, R. L., Basiri, M. L., Kosyk, O., Rossi, M. A., Dichter, G. S., & Stuber, G. D. (2020). Social Stimuli Induce Activation of Oxytocin Neurons Within the Paraventricular Nucleus of the Hypothalamus to Promote Social Behavior in Male Mice. The Journal of Neuroscience, 40(11), 2282. https://doi.org/10.1523/JNEUROSCI.1515-18.2020

Roland, B. L., & Sawchenko, P. E. (1993). Local origins of some GABAergic projections to the paraventricular and supraoptic nuclei of the hypothalamus in the rat [https://doi.org/10.1002/cne.903320109]. Journal of Comparative Neurology, 332(1), 123-143. https://doi.org/10.1002/cne.903320109

Ross, H. E., & Young, L. J. (2009). Oxytocin and the neural mechanisms regulating social cognition and affiliative behavior. Frontiers in neuroendocrinology, 30(4), 534–547. https://doi.org/10.1016/j.yfrne.2009.05.004

Rupp, A. C., Ren, M., Altimus, C. M., Fernandez, D. C., Richardson, M., Turek, F., Hattar, S., & Schmidt, T. M. (2019). Distinct ipRGC subpopulations mediate light’s acute and circadian effects on body temperature and sleep. eLife, 8, e44358. https://doi.org/10.7554/eLife.44358

Tam, S. K. E., Hasan, S., Hughes, S., Hankins, M. W., Foster, R. G., Bannerman, D. M., & Peirson, S. N. (2016). Modulation of recognition memory performance by light requires both melanopsin and classical photoreceptors. Proceedings of the Royal Society B: Biological Sciences, 283(1845), 20162275. https://doi.org/doi:10.1098/rspb.2016.2275

Tang, Y., Benusiglio, D., Lefevre, A., Hilfiger, L., Althammer, F., Bludau, A., Hagiwara, D., Baudon, A., Darbon, P., Schimmer, J., Kirchner, M. K., Roy, R. K., Wang, S., Eliava, M., Wagner, S., Oberhuber, M., Conzelmann, K. K., Schwarz, M., Stern, J. E., … Grinevich, V. (2020). Social touch promotes interfemale communication via activation of parvocellular oxytocin neurons. Nature Neuroscience, 23(9), 1125–1137. https://doi.org/10.1038/s41593-020-0674-y

Theodosis, D. T., Paut, L., & Tappaz, M. L. (1986). Immunocytochemical analysis of the GABAergic innervation of oxytocin- and vasopressin-secreting neurons in the rat supraoptic nucleus. Neuroscience, 19(1), 207–222. https://doi.org/10.1016/0306-4522(86)90016-3

Valle, F. P. (1970). Effects of strain, sex, and illumination on open-field behavior of rats. The American journal of psychology, 103–111.

Xiao, L., Priest, M. F., Nasenbeny, J., Lu, T., & Kozorovitskiy, Y. (2017). Biased Oxytocinergic Modulation of Midbrain Dopamine Systems. Neuron, 95(2), 368–384 e365. https://doi.org/10.1016/j.neuron.2017.06.003

Yu, H., Miao, W., Ji, E., Huang, S., Jin, S., Zhu, X., Liu, M.-Z., Sun, Y.-G., Xu, F., & Yu, X. (2022). Social touch-like tactile stimulation activates a tachykinin 1-oxytocin pathway to promote social interactions. Neuron. https://doi.org/10.1016/j.neuron.2021.12.022

Zhang, Z., Beier, C., Weil, T., & Hattar, S. (2021). The retinal ipRGC-preoptic circuit mediates the acute effect of light on sleep. Nature Communications, 12(1), 5115. https://doi.org/10.1038/s41467-021-25378-w

