## Extended Figures and Legends for "Environmental luminance impairs socio-sexual recognition memory through a succinct retina to supraoptic nucleus circuit"

**Extended Figure 1. Acute light exposure does not significantly alter social memory between a pair of male mice.**

(A) Schematics of pre-social light exposure and the two-trial social recognition test with two male mice.

(B) The investigation duration and recognition index of L and D group ($p = 0.5476$, Mann-Whitney test).

(C) Schematics of pre-social light exposure and the two-trial social recognition test with each trial under total darkness.

(D) The investigation duration and recognition index of L and D group ($*p = 0.0379$, Mann-Whitney test).

**Extended Figure 2. Acute light exposure does not significantly alter sociability in mice.**

(A) Schematics of light exposure and three-chamber test testing sociability (phase 1, P1) and social novelty (phase 2, P2).

(B) Heat map of time spent at different locations during the P1 and P2 stage of three-chamber test for L and D group.

(C) Investigation time spent on the empty and stranger female chamber in L and D group during P1

(D) Investigation time spent on the familiar female and novel female chamber in L and D group during P2

**Extended Figure 3. Acute light exposure does not alter object recognition memory in mice.**

(A) Schematics of light exposure and novel object recognition test (NRT).

(B) Total travel distance and immobile time show no significant difference between the L and D groups.

(C) Interaction time on the object 1 and 2 during the trial 1 of NRT from the L and D groups.

(D) Interaction time of the old object and the new object during the trial 2 of the NRT from the L and D groups

(E) There is no significant difference between the L and D groups for the discrimination index.

**Extended Figure 4. Elimination of M1 or Brn3b$^+$ ipRGCs blocked light-induced**

**c-fos suppression in the SON$^{OT}$ neurons.**

**(A-B)** Representative image of SON$^{OT}$ c-fos expression in the light (A) or dark control (B) group of OPN4$^{DTA/DTA}$ mice. White arrows indicated oxytocin neurons co-labeled with cFos. Scale bar = 50 um.

**(C)** Activation ratio of SON$^{OT}$ neurons in the L and D group of OPN4$^{DTA/DTA}$ mice. There is no significant difference between the L and D groups. $p > 0.9999$, Mann-Whitney test.

**(D-E)** Representative image of SON$^{OT}$ c-fos expression in the light (D) or dark control (E) group of OPN4$^{Cre/+}$; Brn3b$^{z\text{-}DTA/+}$ mice. White arrows indicated oxytocin neurons co-labeled with cFos. Scale bar = 50 um.

**(F)** Activation ratio of SON$^{OT}$ neurons in the L and D group of OPN4$^{Cre/+}$; Brn3b$^{z\text{-}DTA/+}$ mice. There is no significant difference between the L and D groups. $p = 0.6286$, Mann-Whitney test.

**Extended Figure 5. ipRGC forms putative synaptic connects with GABAergic neurons in the pSON.**

(A) Schematics of labeling ipRGC terminals with CTB-Alexa 568 in GAD67-eGFP mice.
(B) Representative confocal image of the pSON region stained with synaptophysin (blue), CTB-Alexa 568 (red), and GFP from GABAergic neurons (green). Scale bar = 10 um.
(C) Enlarged image of a GAD67 positive cell with colocalization of CTB-Alexa 568 and synaptophysin. White color indicates triple colocalization.
(D-F) Verification of colocalization at different orthogonal sections.

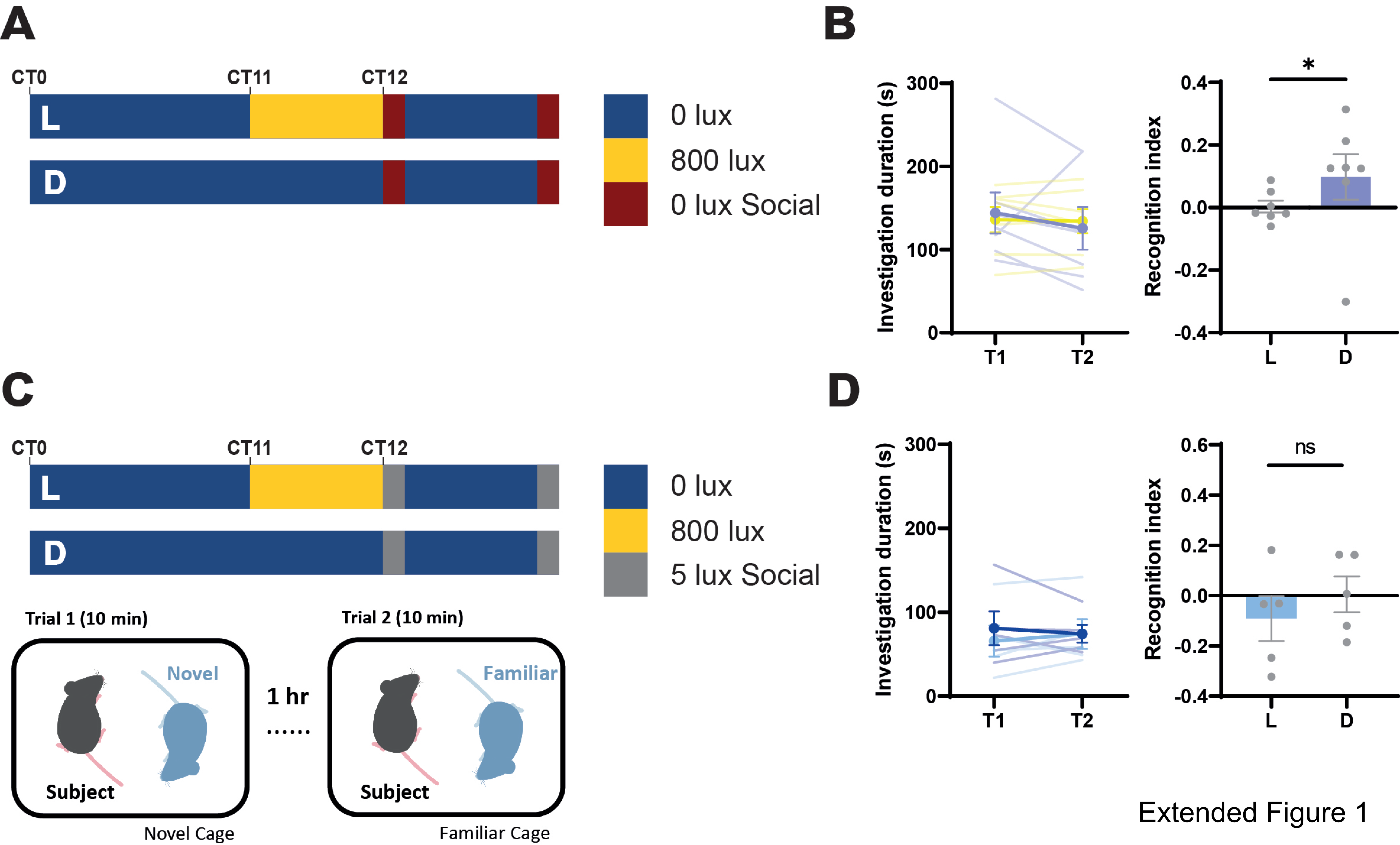


Extended Figure 1

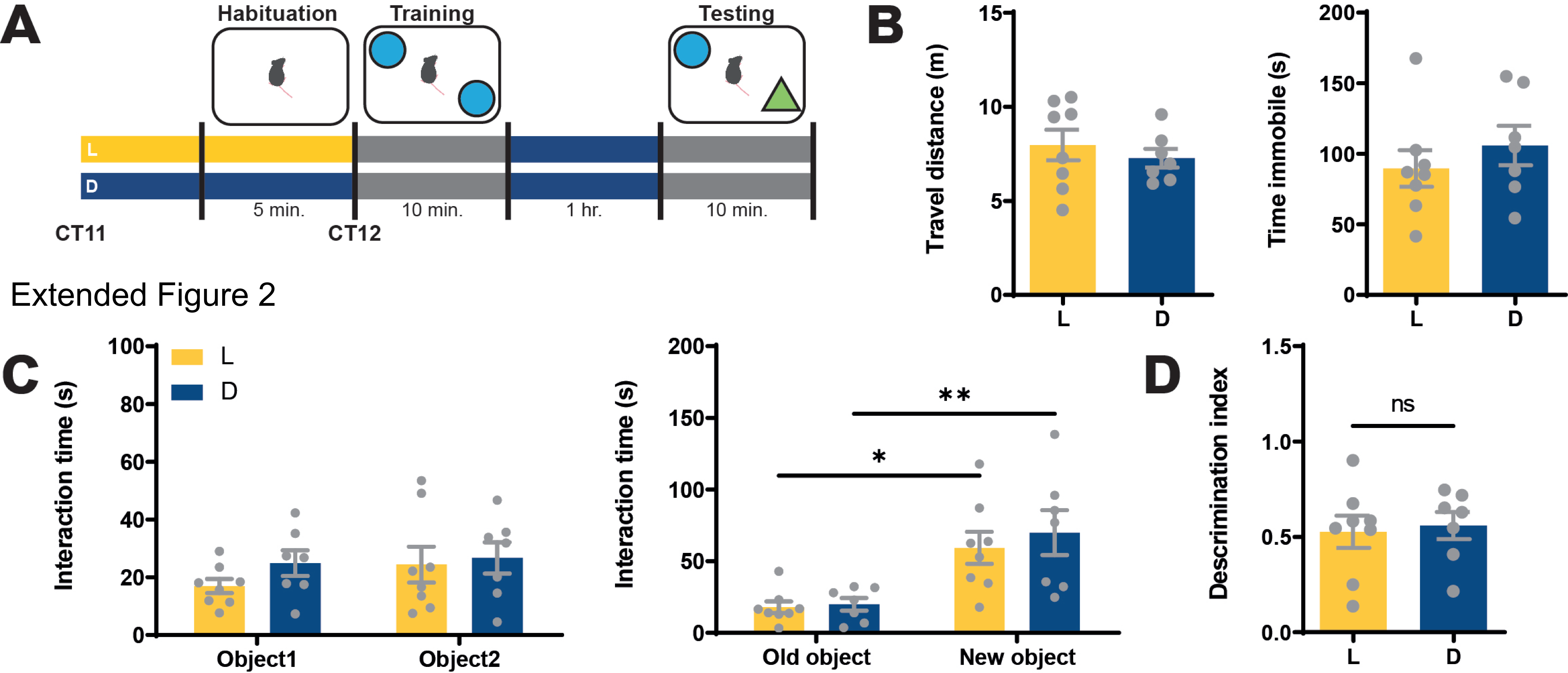


Extended Figure 2

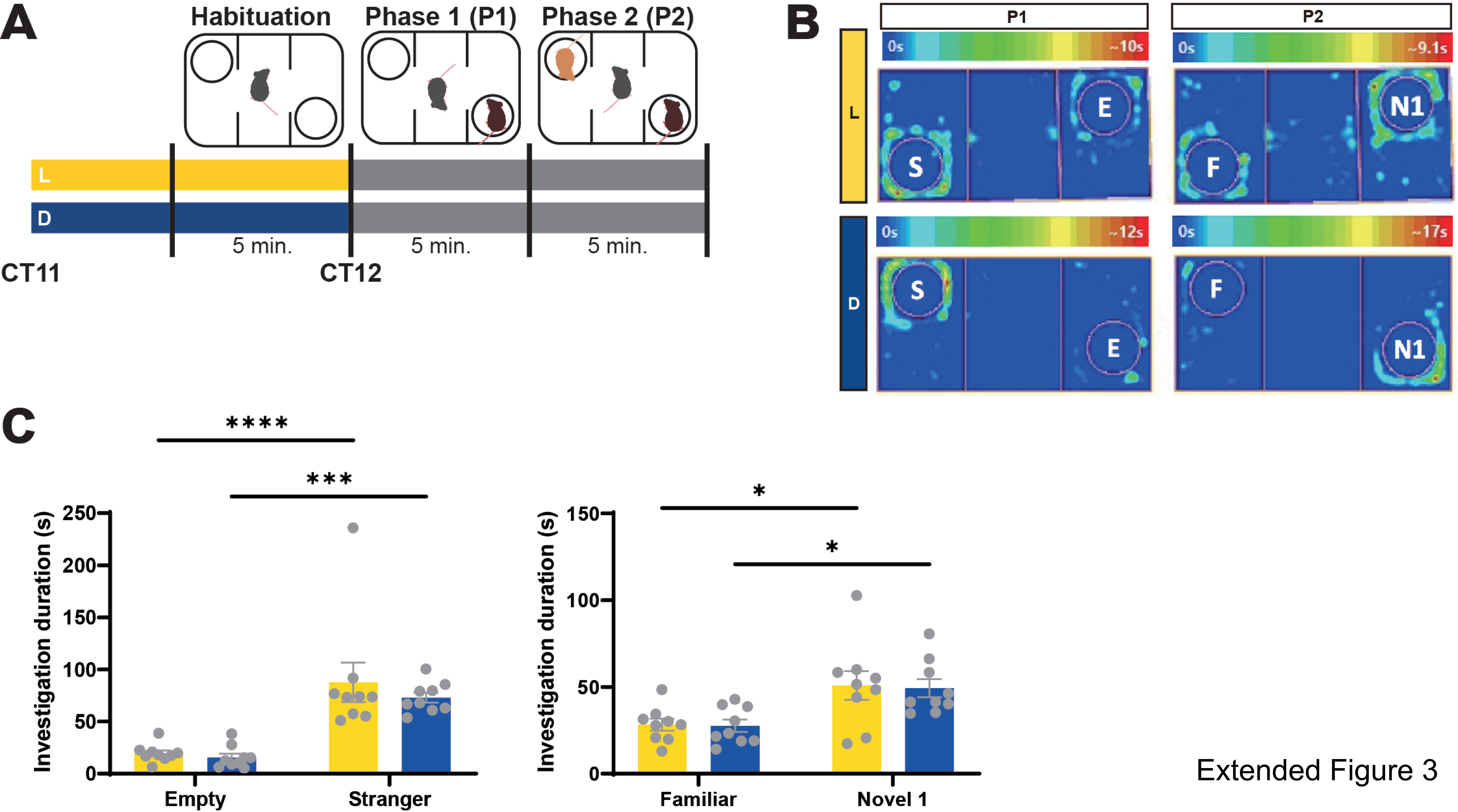


Extended Figure 3

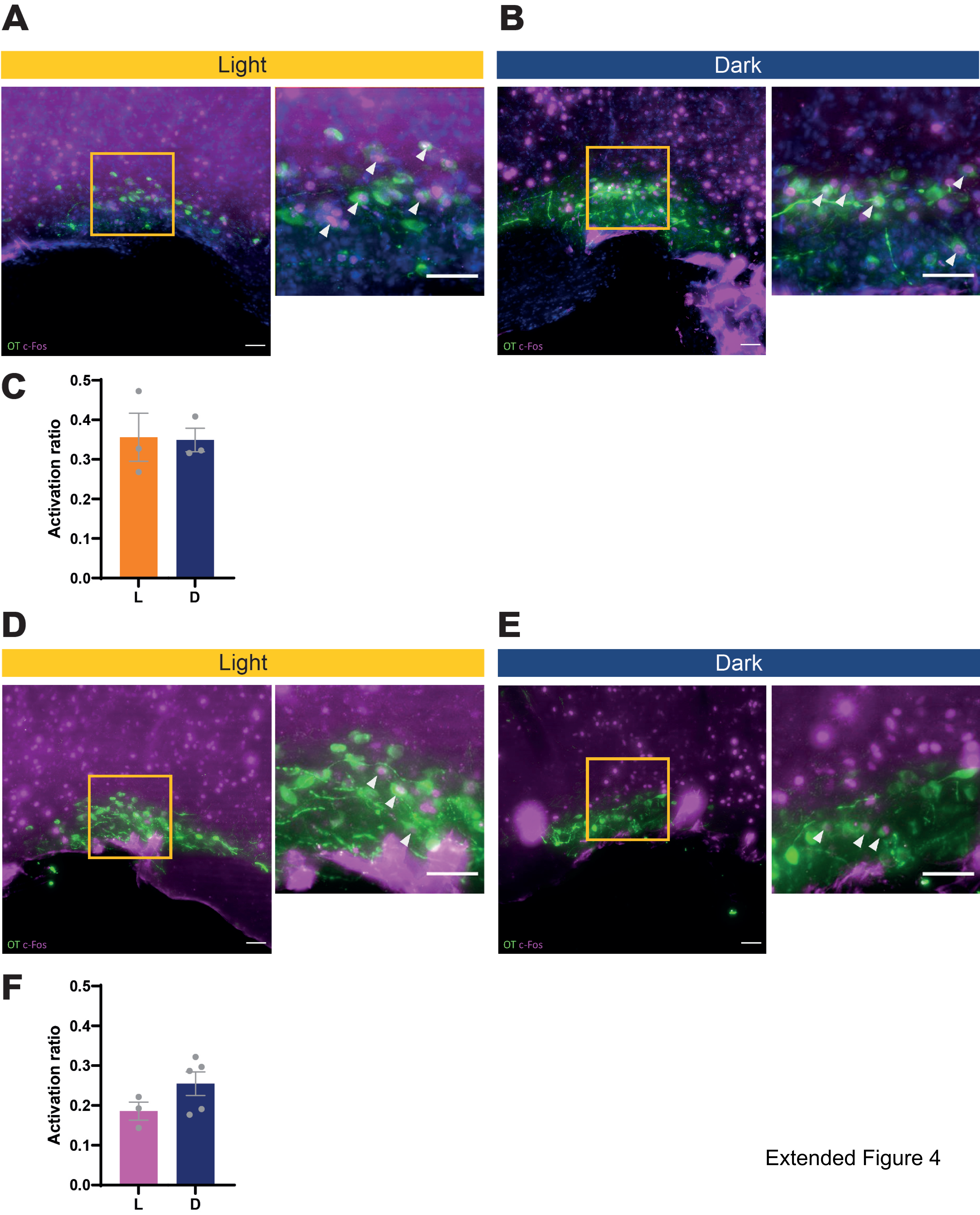


Extended Figure 4

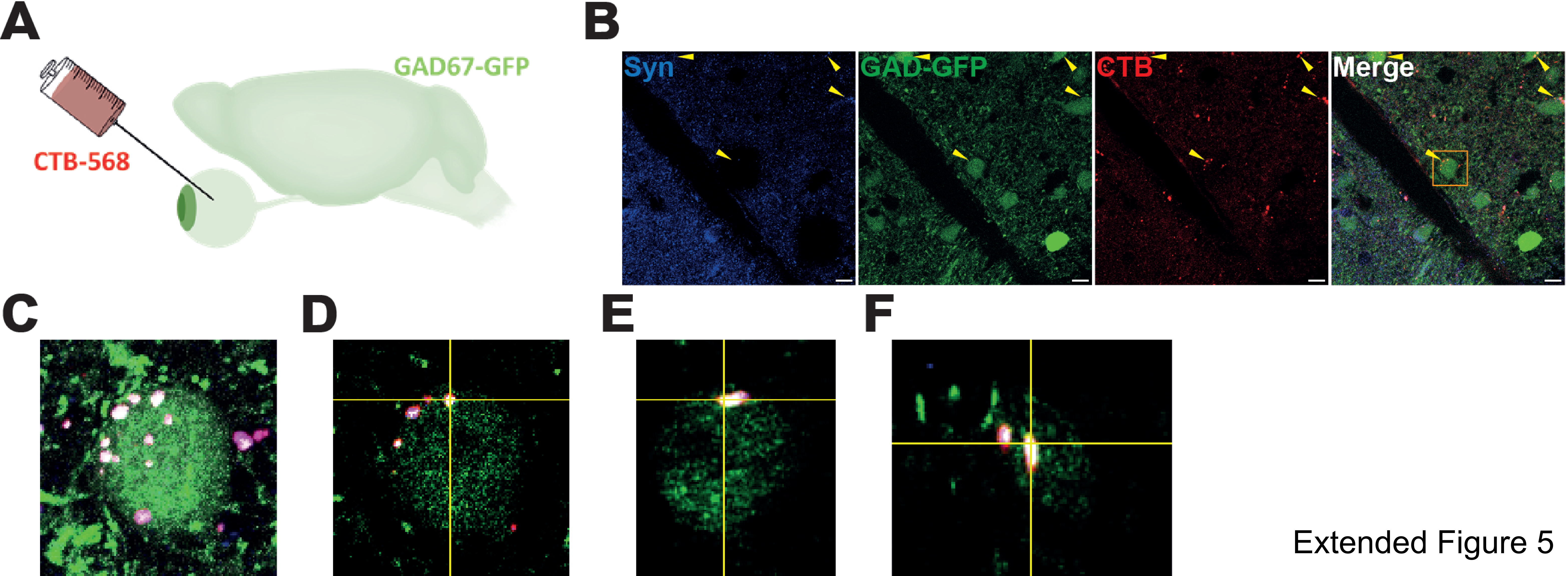


Extended Figure 5
